# From Easy to Hopeless - Predicting the Difficulty of Phylogenetic Analyses

**DOI:** 10.1101/2022.06.20.496790

**Authors:** Julia Haag, Dimitri Höhler, Ben Bettisworth, Alexandros Stamatakis

## Abstract

Phylogenetic analyses under the Maximum Likelihood model are time and resource intensive. To adequately capture the vastness of tree space, one needs to infer multiple independent trees. On some datasets, multiple tree inferences converge to similar tree topologies, on others to multiple, topologically highly distinct yet statistically indistinguishable topologies. At present, no method exists to quantify and predict this behavior. We introduce a method to quantify the degree of difficulty for analyzing a dataset and present Pythia, a Random Forest Regressor that accurately predicts this difficulty. Pythia predicts the degree of difficulty of analyzing a dataset *prior* to initiating Maximum Likelihood based tree inferences. Pythia can be used to increase user awareness with respect to the amount of signal and uncertainty to be expected in phylogenetic analyses, and hence inform an appropriate (post-)analysis setup. Further, it can be used to select appropriate search algorithms for easy-, intermediate-, and hard-to-analyze datasets.

## Introduction

The goal of a phylogenetic inference is to find the phylogenetic tree that best explains the given biological sequence data. Since the number of possible tree topologies grows super-exponentially with the number of taxa, one cannot compute and score every possible tree topology. Instead, one deploys tree inference heuristics that explore the tree space to find a tree with a ‘good’ score, for example under the Maximum Likelihood (ML) criterion (Yang *et al*. 1995). However, these heuristics do not guarantee that the tree inference will converge to the *globally* optimal tree. Therefore, under ML, one typically infers multiple trees and subsequently summarizes the inferred, *locally* optimal trees via a consensus tree. One can observe that for some datasets, all individual, independent ML tree searches converge to topologically similar trees. This suggests that the likelihood surface of such datasets exhibits a single likelihood peak, yielding the dataset easy to analyze. For other datasets, one observes that the independent tree inferences converge to multiple topologically distinct, yet, with respect to their ML score, statistically indistinguishable, locally optimal trees. These datasets are hence difficult to analyze, and we say that they exhibit a rugged likelihood surface. This diverse behavior of phylogenetic tree searches has already been reported in several publications (Lakner *et al*. 2008; Stamatakis 2011; Morel *et al*. 2020). In general, the more tree inferences we perform, the better our understanding of the dataset’s behavior and coverage of the respective tree space will be. However, under ML, inferring a single tree can already require multiple hours or even days of CPU time. In order to save time and resources, an optimal analysis setup will perform as few tree inferences as necessary. For easy-to-analyze datasets with a single likelihood peak, we require fewer and less involved tree search heuristics and bootstrap replicate searches to adequately sample the tree space, as opposed to difficult-to-analyze datasets with rugged likelihood surfaces. To the best of our knowledge, and despite anecdotal reports on the behavior of difficult datasets, there does not yet exist a *quantifiable* definition of dataset difficulty that captures the behavior of ML tree searches on datasets.

In order to speedup ML tree inferences, researchers have developed elaborate ML tree inference tools that combine multiple search strategies to reduce the risk of becoming stuck in local optima. There also exist early-stopping criteria to determine whether the tree inference has converged. Such early-stopping methods deploy ad hoc or statistical criteria to terminate the tree inference. For example, the ML tree inference software FastTree (Price *et al*. 2010) relies on a maximum number of topology optimization iterations as a function of the number of sequences in the dataset. The ML software RAxML (Stamatakis 2014) implements an early-stopping criterion based on the topological distance between the respective best trees found in two consecutive optimization cycles (Stamatakis 2011). Vinh and von Haeseler (2004) propose an estimation criterion that determines with 95% confidence whether continuing the tree inference will yield a better tree than the currently best tree. However, early stopping criteria only determine the convergence of the current tree search, but they do evidently not guarantee that the search has converged to the globally optimal tree. Thus, to better characterize and explore the tree search space, additional tree inferences and subsequent a posteriori analyses are required. In contrast, assessing the expected behavior of a dataset prior to conducting compute-intensive tree inferences allows for a more informed decision on the most appropriate tree inference and post-analysis setup. It also allows users to reassemble/modify difficult datasets as these will most likely require resource-intensive analyses that yield contradicting, yet almost equally likely, tree topologies with low confidence. Several methods have already been developed to assess the information content of datasets *prior* to tree inference, the most prominent example being the treelikeness of a dataset (Bandelt and Dress 1992; Lyons-Weiler *et al*. 1996; White *et al*. 2007). Simple and fast-to-compute metrics include the sites-over-taxa ratio. For instance, Rosenberg and Kumar (2001) conclude that a higher phylogenetic inference accuracy can be achieved by increasing the MSA length, rather than including more taxa/sequences. A more involved method was proposed by Holland *et al*. (2002). The authors suggest the use of *δ*-plots, that is histograms, based on all quartet distances in the Multiple Sequence Alignment (MSA). However, computing the *δ*-plots is time-intensive due to the computational complexity of O(*n*^4^), where *n* is the number of taxa in the MSA. Misof *et al*. (2014) provide an overview of various methods for calculating the treelikeness, prior to a phylogenetic analysis. The authors acknowledge that the considered treelikeness estimation methods capture certain aspects of the MSAs. However, they conclude that none of them sufficiently informs the user about the expected behavior of phylogenetic analyses in general, and suggest further research in this area.

### New Approach

Here, we initially introduce a quantification of difficulty based on the result of 100 ML tree inferences per MSA. We then show that this quantification adequately represents the behavior of the ML searches on the dataset. Since executing 100 ML tree searches is computationally prohibitive in general, we train a Random Forest Regressor (Ho 1995) that can predict the difficulty of a given MSA that is exclusively based on MSA attributes and some fast and thus substantially less expensive parsimony-based tree inferences (Farris 1970; Fitch 1971). By extracting multiple simple and fast-to-compute attributes, such as the sites-over-taxa ratio, and by deploying machine learning, we devise an accurate difficulty predictor called *Pythia*. We attain a high prediction accuracy, with a mean absolute prediction error (MAE) of 0.09 and a mean absolute percentage error (MAPE) of 2.9 %. Computing the prediction features and predicting the difficulty is on average approximately five times faster than a single ML tree inference. Pythia predicts the difficulty of a dataset on a scale ranging between 0.0 (easy) to 1.0 (difficult).

In contrast to the aforementioned early-stopping criteria that can be applied during ML searches, Pythia informs the user about the expected behavior of the MSA in ML phylogenetic analysis *prior* to any ML phylogenetic inference. Thereby, users can take informed decisions on the most appropriate ML analysis and post-analysis setup. This includes, for example, a careful consideration of the number of required independent, resource-intensive, tree searches based on the difficulty. Also, for difficult MSAs, the user will be able to improve the informativeness of the MSA, for example by increasing sequence length or removing sequences, to assemble an MSA that is easier to analyze.

Thereby, one can save valuable time and resources by not performing tree inferences on difficult MSAs. We therefore suggest that an analysis with Pythia should be conducted at the beginning of any ML phylogenetic analysis. Note that the predicted difficulty does not directly predict the number of tree inferences required to sufficiently sample the tree space, as this number also depends on the implemented tree inference heuristic.

Pythia is available as open source software libraries in C and Python. Both libraries include the trained Random Forest Regressor and the computation of the required prediction features. The C library CPythia is an addition to the COre RAXml LIBrary (Coraxlib) (Exelixis-Lab 2022) and is available at https://github.com/tschuelia/CPythia. Additionally, we provide PyPythia, a lightweight, stand-alone Python library, including a respective command line interface. PyPythia is available at https://github.com/tschuelia/PyPythia. Finally, by using the phylogenetic tree data that is being collected by our dynamically growing RAxML Grove (Höler *et al*. 2021) database, we regularly retrain Pythia and update the predictor in both libraries.

## Results

### Difficulty Prediction Accuracy

Our training data contains 3250 empirical MSAs obtained from TreeBASE (Piel *et al*. 2000). We divide this training data into a *training set* (80 %) and a *test set* (20 %). The training set is used for training the predictor and the test set is exclusively used for evaluating the trained predictor. Pythia predicts the degree of difficulty on a scale between 0.0 to 1.0. A value of 1.0 indicates a difficult (hopeless) MSA with a rugged tree space. We expect such an MSA to exhibit multiple, statistically indistinguishable locally optimal yet topologically highly distinct trees. In contrast, we expect an MSA with a value of 0.0 to be easy to analyze by requiring only few independent tree searches. Pythia attains a mean absolute error (MAE) of 0.09. This corresponds to a mean average percentage error (MAPE) of 2.9 %. The mean squared error (MSE) is 0.02 and the R^2^ score is 0.79. Supplementary Figures S5a and S5b show the distribution of prediction errors for the training data. When analyzing the prediction error, we notice that Pythia tends to overestimate the difficulty of MSAs with a difficulty ≤ 0.3 and underestimate the difficulty for MSAs with a difficulty > 0.3 (Supplementary Figure S4). We suspect that this is caused by an uneven distribution of difficulties in the training data. Our training data contains substantially more ‘easy’ MSAs than difficult MSAs: for approximately 60 % of MSAs the assigned difficulty is ≤ 0.3 and only about 10 % have a difficulty ≥ 0.7 (Supplementary Figure S2).

### Feature Importance

In our study, we analyze a plethora of distinct features of the MSA, of trees inferred under parsimony, and features based on a single ML tree inference using RAxML-NG. In order to decrease the runtime of Pythia’s difficulty prediction, we analyze the runtime of computing each feature for all MSAs in our training data, as well as the importance of the feature for the prediction. Based on these results, we selected a subset of eight features:

- Sites-over-taxa ratio:

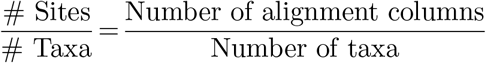
- Patterns-over-taxa ratio:

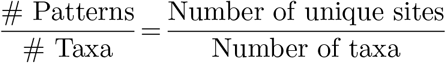
- % Invariant sites: Percentage of fully conserved sites.
- % Gaps: Proportion of gaps in the MSA.
- Entropy: Shannon Entropy (Shannon 1948) as average over all per-column/site entropies. See the supplementary information for a more detailed description.
- Bollback Multinomial: Multinomial test statistic according to Bollback (Bollback 2002). See the supplementary information for a more detailed description.
- RF-Distance Parsimony Trees: RF-Distances between 100 trees inferred using parsimony.
- % Unique Topologies Parsimony Trees: Percentage of unique topologies among the 100 inferred parsimony trees.

Four of these are direct attributes of the MSA: the sites-over-taxa ratio, the patterns-over-taxa ratio, the percentage of gaps, and the percentage of invariant sites. Two features quantify the amount of information in the MSA: the Shannon entropy (Shannon 1948) and the Bollback multinomial (Bollback 2002). Two additional features are based on rapid parsimony tree inferences: we infer 100 parsimony trees via a randomized step-wise addition order procedure and compute their average pair-wise topological distances using the Robinson-Foulds distance metric (RF-Distance) (Robinson and Foulds 1981), as well as the proportion of unique topologies in this set of 100 parsimony trees. In Supplementary Information Section 2, we present all features we considered and analyzed in more detail, alongside the respective feature importance and runtime to justify the selection of the eight features we finally use. Table 1 shows the prediction importances of the eight features upon which the difficulty prediction is based. We use the permutation importance (Breiman 2001) for computing feature importance. As the table shows, the difficulty prediction heavily relies on the average RF-Distance and the proportion of unique topologies among the inferred parsimony trees. This is expected, as our difficulty definition under ML reflects the ruggedness of the tree space and correlates well with the ruggedness under parsimony.

**Table 1.** Importance of the subset of features we use to train Pythia.

| Feature | Impurity importance |
| --- | --- |
| % Unique topologies<br>parsimony trees | 42.9 % |
| RF-Distance<br>parsimony trees | 33.2 % |
| Entropy | 17.0 % |
| Patterns-over-taxa | 13.6 % |
| % Gaps | 2.5 % |
| Bollback | 2.3 % |
| Sites-over-taxa | 1.5 % |
| % Invariant | 0.6 % |

### Runtime of Feature Computation

Computing the selected set of prediction features takes on average 5 ± 31 s (*μ*±*σ*) with a median runtime of 1 s. For our training data, this corresponds to a runtime of 21.5 ± 88.6 % relative to the runtime for inferring a single ML tree using RAxML-NG. The median is 6.8 %. The high average compared to the median, and the large spread, are due to the fact that the runtime of computing the prediction features predominantly depends on the size of the MSA. The larger the MSA, the faster the feature computation is compared to a single ML tree inference. Supplementary Figure S3 depicts this correlation. For benchmarking the runtimes of the feature computation, we used the implementation in our Python library. When running a subsequent ML tree inference, the runtime overhead induced by the prediction can be amortized by passing the inferred maximum parsimony trees as starting trees to the ML inference tool (e.g. RAxML-NG). Instead of re-computing parsimony starting trees, the RAxML-NG simply initiates its tree searches on the provided parsimony starting trees.

## Discussion

Predicting the difficulty of MSAs to gain a priori insights into the expected behavior of phylogenetic tree searches and the shape of the likelihood surface constitutes a vital step towards faster phylogenetic inference and a more targeted setup of the computational analyses and post-analyses. Our difficulty prediction allows for careful consideration of the number of tree inference required to sufficiently sample tree space *prior* to ML analyses. Especially for easy MSAs, this has the potential to save valuable time and resources. In this paper, we presented a quantifiable definition of difficulty for MSAs and showed that this definition adequately represents the ruggedness of the tree space of the dataset under ML. Using this definition, we trained Pythia, a Random Forest Regressor, to predict the difficulty on a scale ranging between 0.0 to 1.0. We showed that Pythia achieves a high prediction accuracy. We further showed that the runtime to compute the prediction features is on average only approximately one fifth of the runtime required for inferring a single ML tree with RAxML-NG. The more taxa and sites the MSA has, the faster the feature computation is relative to a single ML tree inference, making Pythia especially valuable for phylogenetic analyses on MSAs with many sites and taxa. We conclude that predicting the difficulty of an MSA prior to any tree inference allows for faster analyses, informing user expectations regarding the stability of the inferred tree, and Pythia should be included in ML phylogenetic inference pipelines. As a cautionary note, we emphasize that the ruggedness of the tree space might also depend on the model and tree inference heuristic being used. Yet, the fact that Pythia relies on parsimony trees to predict the ruggedness of ML trees shows that there exists a correlation between models regarding the ruggedness of the tree space and thus, the difficulty of the analysis.

Using our dynamically growing RAxML Grove database, we perpetually enlarge our training data and retrain Pythia at regular intervals. The goal of this retraining is to continuously improve the predictive power of Pythia by providing more, and more diverse data in terms of the distribution of feature values. At the time of writing this paper, the difficulty labels in our training data are unevenly distributed. Since we carefully select the new MSAs from RAxML Grove we include for retraining (see Section Retraining the Model), we expect the effect of uneven label distribution to cancel out over time.

### Use and Misuse of Pythia

We suggest predicting the difficulty using Pythia prior to any ML phylogenetic inference, as this will allow for more targeted analysis setups. For example, for a difficult MSA, the user should be careful to report a single ML tree as best-known tree, as the tree space most likely exhibits multiple, indistinguishable local optima. The user should also be aware that a more difficult MSA requires a higher number of independent tree searches to construct a reliable consensus tree. Furthermore, difficult MSAs require a more careful consideration of necessary additional phylogenetic analyses and post-processing steps. Especially for very difficult MSAs (difficulty > 0.8) we suggest to consider improving upon the difficulty of the MSA prior to analysis. This is because a phylogenetic analysis on very difficult MSAs, will most likely not yield a well-resolved tree, even if a consensus of numerous almost equally likely yet topologically distinct ML trees is built. Pythia is not intended to directly predict the number of independent tree searches required for conducting a thorough ML analysis, as this number also heavily depends on the search strategy of the respective ML inference tool.

### Future Work

Potential future applications of Pythia include, for instance, the assembly of benchmark datasets which cover a broad and representative difficulty range for testing novel phylogenetic models and tools. Pythia can also serve as a criterion during the empirical dataset assembly process. For instance, additional sequence data can be added to yield a dataset that is easier to analyze.

Another avenue for future work is to implement a difficulty-aware tree inference heuristic. Depending on the difficulty of the MSA, we can, for example, apply different heuristic search strategies. For instance, on easy MSAs it might be sufficient to explore the tree space via a less thorough exploration strategy, that is, by only using Nearest-Neighbor-Interchange (NNI) moves. In comparison to Subtree Pruning and Regrafting (SPR) moves, this reduces the tree topology search complexity from 𝒪 (*n*^2^) to 𝒪 (*n*) (Heath and Ramakrishnan 2010).

In our study, we focused on predicting the difficulty of ML phylogenetic inferences. Another popular method to explore the tree space of an MSA is Markov chain Monte Carlo (MCMC) based Bayesian phylogenetic inference. Since both methods, ML and MCMC, rely on the same input MSA and on the same likelihood function, we suspect the difficulty to be reflected in the apparent convergence speed of MCMC methods. In this section, we will explore this potential correlation on three exemplary MSAs.

Besides informing the computational setup of ML phylogenetic analyses, Pythia can also potentially be applied to adjust user expectations regarding the bootstrap support of the best-known tree as well as related support measures. For instance, the perhaps most common and recurrent user inquiry on the RAxML Google user support group concerns possible reasons for often unexpected and disappointingly low bootstrap support values. In this section, we also present an exploratory analysis of the correlation between the difficulty as predicted by Pythia, and the bootstrap support values for three MSAs.

Since both, MCMC phylogenetic analyses and bootstrap analyses, constitute extremely time- and resource-intensive tasks, a thorough exploration of their connection to difficulty prediction is beyond the scope of this work.

#### MCMC Convergence Prediction

The features we use to predict the difficulty of an MSA are independent of the inference method used for the subsequent analyses. However, as we describe in the Quantification of Difficulty subsection, our difficulty quantification is based on 100 tree inferences using RAxML-NG which implements the ML method. Therefore, our predictions might be biased towards ML analyses and potentially not describe the ruggedness of the tree space in a model-independent manner. To assess if our predictions can be generalized, we compare our difficulty prediction to convergence diagnostics of MCMC based phylogenetic analyses. For three DNA MSAs (D27 (Hedges *et al*. 1990), D125 (Poulakakis and Stamatakis 2010), and D354 (Grimm *et al*. 2006)) we perform MCMC analysis using MrBayes (Ronquist *et al*. 2012). We run four chains for 10 million generations each using the general time reversible (GTR) model with four Γ rate categories to account for among site rate heterogeneity. MrBayes reports the average standard deviation of split frequencies (ASDSF; split frequencies: relative number of occurrence of splits/bipartitions in the set of posterior trees) as a convergence diagnostic metric and suggests executing additional generations as long as the ASDSF is ≥ 0.01. D125 is an easy dataset with an expected clear, single likelihood peak. The difficulty according to our definition is low (≪ 0.1) and MrBayes appears to converge: the ASDSF value drops below 0.01 after 150000 generations and is ≪ 0.01 after only 1 million generations. D27 exhibits at least two distinct likelihood peaks, suggesting that the MSA is rather difficult to analyze (Lakner *et al*. 2008). The difficulty according to our definition is 0.45 and after 10 million generations MrBayes reports an ASDSF of 0.011, indicating that the MCMC did not converge to a single local optimum. D354 exhibits a rugged likelihood surface (Grimm *et al*. 2006), so we expect a high difficulty and no convergence. The assigned difficulty for D354 is 0.6 and after 10 million generations the ASDSF is 0.009. According to MrBayes this suggests convergence and adding more generations should improve the ASDSF. However, we observe that the ASDSF did not improve during the last 2 million generations, and adding more generations did not further improve the ASDSF. D125 with 125 taxa and approximately 30000 sites is a larger dataset than D354 with 354 taxa and only 460 sites. Yet, D125 converges after 1 million generations, while for D354 the ASDSF drops below 0.01 only after 8 million generations. The smallest dataset D27 with 27 taxa and 1940 sites indicates no convergence after 10 million generations according to the ASDSF. We thus suspect that the number of generations required for the MCMC is correlated to the difficulty rather than to the size of the dataset.

#### Bootstrap Support Values

As already mentioned, the perhaps most common question on the RAxML user support Google group is related to disappointingly low support values. We expect the difficulty, and thus the vastness of the tree space, to correlate with the support values of the best-known tree in a subsequent bootstrapping analysis. We use the same MSAs for the same reasons as for the exploratory MCMC convergence prediction conducted above: D27, D125, and D354. For each MSA, we run RAxML-NG using its --all execution mode. This mode infers 20 ML trees for the MSA, infers bootstrap replicate trees, and draws support values on the tree with the highest log-likelihood (best-known tree). Per default, RAxML-NG infers at most 1000 bootstrap replicates, but implements an early-stopping criterion that determines convergence based on the bootstopping criterion presented by Pattengale *et al*. (2010). To explore the correlation between the difficulty prediction value and the bootstrap support values, we compute the average and standard deviation *μ*±*σ* of bootstrap support values on the respective best-known trees. As stated above, D125 is an easy dataset exhibiting a clear signal with an assigned difficulty ≪ 0.1. This is reflected by the high bootstrap support values: *μ*±*σ* = 97.64±8.38 %. The assigned difficulty for D27 is 0.45 and RAxML-NG reports the bootstrap support values as *μ*±*σ* = 51.5±29.02 %. Dataset D354 is the most difficult among the three example MSAs with a predicted difficulty of 0.6. Hence, the bootstrap support values are the lowest among the three MSAs with *μ*±*σ* = 43.41±32.48 %.

## Materials and Methods

We formulate the difficulty prediction challenge as a supervised regression task. The goal is to predict the difficulty on a scale ranging between 0.0 (easy) to 1.0 (difficult). We face two main challenges: (i) obtaining a sufficiently large set of MSAs to train Pythia on, ideally consisting of empirical MSAs, and (ii) obtaining ground-truth difficulties that represent the actual difficulty of the training data. In the following, we present how we obtain the training data and assign ground-truth difficulties. We further present our trained regression model, and finally present our heuristic for regularly retraining the regression model to continuously improve the prediction accuracy of Pythia.

### Quantification of Difficulty

In order to train a reliable difficulty predictor, we need a reliable ground-truth label for each training datum. To obtain such labels, we require a quantifiable difficulty definition. To stringently quantify the difficulty of an MSA, we would have to explore the entire tree space. Since this is computationally not feasible, we need to rely on a heuristic definition. Our heuristic to quantify the difficulty is based on 100 ML tree inferences. In our analyses, we use RAxML-NG. First, we infer *N*_all_ = 100 ML trees and compute the average pairwise relative RF-Distance between all trees (*RF*_all_), as well as the number of unique topologies among the 100 inferred trees 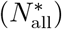.

We determine the best tree among the 100 inferred trees according to the log-likelihood, and compare all trees to this best tree using statistical significance tests. We assign trees that are not significantly worse than the best tree to a so-called *plausible tree set*. In our analyses, we use the statistical significance tests as implemented in the IQ-TREE software package (Minh *et al*. 2020). Due to the continuing debate about the most appropriate significance test for comparing phylogenetic trees, we use the approach suggested by Morel *et al*. (2020): we only include trees that pass *all* significance tests in the plausible tree set. We further refer to the number of trees in this plausible tree set as *N*_pl_. We compute the average pairwise relative RF-Distances between trees in the plausible tree set (*RF*_pl_), as well as the number of unique topologies 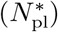. Finally, we compute the difficulty of the dataset based on the following formula:

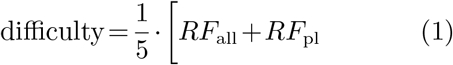

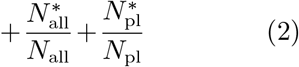

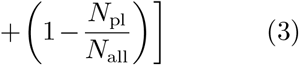

The reasoning for expression (1) is that if the RF-Distance is high, the tree space consists of multiple distinct, locally optimal tree topologies which characterize a dataset that is difficult to analyze. With expression (2) the reasoning is that the tree surface becomes more rugged, the more distinct locally optimal tree topologies the tree inference yields, and the more tree topologies are not significantly different from the best tree. Finally, the rationale for expression (3) is that, the more tree inferences yield a plausible tree, the more informative the MSA will be about the underlying evolutionary process and the easier the MSA will be to analyze. Each term is a value between 0.0 and 1.0, leading to an average value between 0.0 and 1.0 that quantifies the overall difficulty.

For each MSA in our training data, we compute the difficulty according to this definition. To this end, we implement a training data generation pipeline that automatically performs all required tree inferences, statistical tests, and computes the difficulty label alongside the features required for training Pythia. We implement this pipeline using the Snakemake workflow management system (Köster and Rahmann 2012) and Python 3. The pipeline code is available at https://github.com/tschuelia/difficulty-prediction-training-data. In Supplementary Section 6 we list the software versions we use in the described pipeline.

### Training Data

We train Pythia using empirical MSAs obtained from TreeBASE (Piel *et al*. 2000). To date, our training data consists of 3250 MSAs, of which 74% contain DNA data and 26% contain Amino Acid (AA) data. The training data includes partitioned and unpartitioned MSAs. We provide a detailed overview of the training data in Supplementary Section 1. We include DNA and AA data in the same setup as, according to our analyses, the prediction behaves analogously on both data types. We provide a more thorough justification of this equal treatment of DNA and AA data in Supplementary Section 5. Note that while we include partitioned MSAs in our training data, we compute all features across the entire MSA regardless of the defined partitions. The high feature importance of the parsimony tree based features, as well as the entropy that are all partition-agnostic, justifies this choice.

Figure 1 depicts the workflow for training data generation. For each MSA, we compute the difficulty according to the above definition as ground-truth label for supervised training using the training data generation pipeline. We compute the corresponding prediction features using our Python library. The set of prediction features and the corresponding difficulty label form our training data. For training the regression model, we split this training data into two sets: a training set and a test set. The training set comprises 80% of the training data and the test set the remaining 20%. The test set is exclusively used for evaluating the predictive power of the difficulty predictor. To ensure an even distribution of difficulty labels in the training and test sets, we deploy stratified sampling. Stratified sampling splits all difficulty labels into disjoint subsets and draws random samples from each subset independently. In principle, using simulated data would allow us to increase the size of the training data. However, since simulating data that behaves analogously to empirical data under ML tree inferences constitutes a challenging task (Höhler *et al*. 2021), we decided against using any simulated data.

**FIG. 1.**
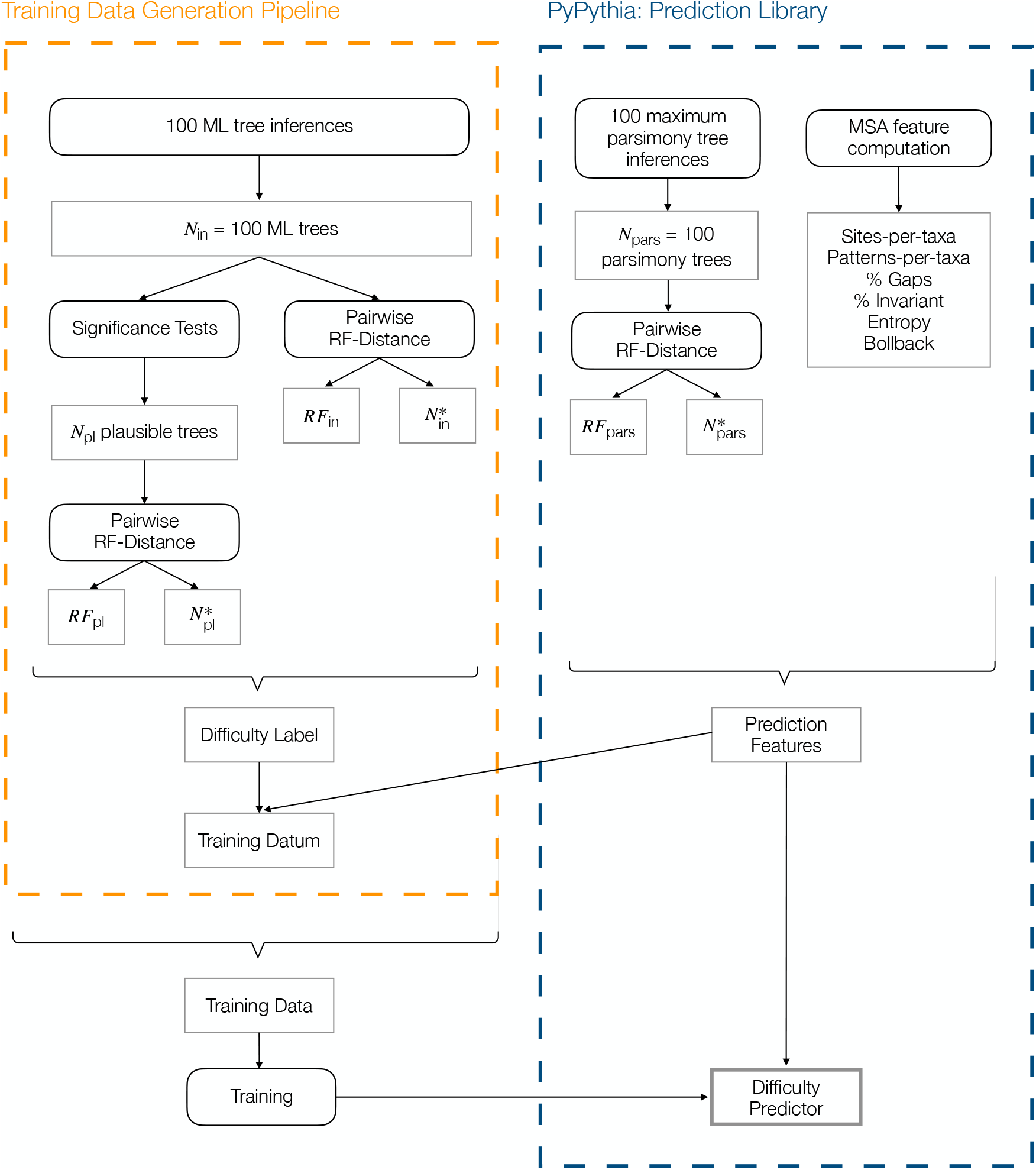
Schematic depiction of the training data generation procedure. For each MSA, we compute the difficulty label based on our difficulty quantification using our training data generation pipeline (left dashed box). We further compute the prediction features using our Python prediction library PyPythia (right dashed box). Using the difficulty label and the corresponding prediction features for all MSAs in our training data, we train Pythia.

### Label Validation

Due to the lack of absolute ground-truth labels, we need to rely on the inferred difficulty labels. The motivation of the difficulty prediction is to limit the number of tree inferences required to sufficiently sample the tree space and obtain a representative consensus tree. To verify the label assignment for each dataset, we conduct two analyses. First, we compare the consensus tree obtained from the plausible tree set constructed from all 100 ML tree inferences (*baseline tree*) to the consensus of the plausible trees we obtain when inferring only 100 * *difficulty* trees (*prediction tree*). Note that for this analysis we use the *difficulty* we compute according to the above definition rather than using a predicted difficulty. We compare the topologies of the consensus trees using the RF-Distance. The RF-Distance between the *baseline tree* and the *prediction tree* is on average 9.6±15.8 %. This noticeable topological difference suggests that either a) the difficulty labels do not sufficiently represent the tree search behavior of the dataset, or b) 100 tree inferences do not sufficiently sample the tree space. To determine the impact of b), we repeatedly sample 99 trees out of the 100 tree inferences and compute the consensus tree *C*_*i*_ of the respective plausible tree set. We then assess the average RF-Distance between all consensus trees *C*_*i*_. For our training data, this RF-Distance is on average 8.1±14.5 %. We conclude that mostly b) causes the high topological distances between the *baseline tree* and the *prediction tree*. In fact, a high RF-Distance between the consensus trees *C*_*i*_ for an MSA is correlated with its difficulty. The Spearman’s rank correlation coefficient is 0.88 with a p-value of 0.0 (≪ 10^−300^). Thus, the more difficult the MSA, the higher the topological distances between the consensus trees *C*_*i*_ will be. The second analysis to justify our quantification of difficulty ensures that selecting the number of tree inferences based on the difficulty does not negatively impact the quality of the tree inference. As stated above, the difficulty can, in general, not predict the number of tree searches required to sufficiently sample the tree space, as this number also depends on the implemented tree inference heuristic. However, since we define the difficulty based on 100 ML tree inference in RAxML-NG, we can use the difficulty to determine the number of required tree inferences when again using RAxML-NG as a fraction of 100. Thus, to analyze the influence of the difficulty on the quality of the tree inference, we compare the log-likelihoods obtained from 100 independent RAxML-NG tree searches (*LnLs*_100_) to the log-likelihoods of |*difficulty* ·100| tree searches (*LnLs*_diff_) for all MSAs in our training data. We compare the respective best found log-likelihoods 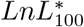 and 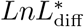, as well as the average log-likelihoods 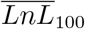 and 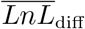.

For 81 % of the MSAs, the best found log-likelihoods 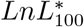 and 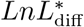 are identical. For the remaining 19 % of MSAs, 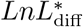 is on average ≪0.01 % worse than 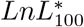. The average log-likelihoods 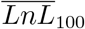 and 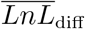 deviate on average by 0.01 % only.

This analysis only serves for justifying the definition of our difficulty quantification. Predicting the number of tree inferences as a fraction of 100 is only applicable to ML tree inference with RAxML-NG. It should further be mentioned, that RAxML-NG infers only 20 trees by default and simply increasing the number of tree inferences to |*difficulty* ·100| is discouraged.

Given these analyses, we conclude that our difficulty quantification is sufficiently accurate to capture the tree search complexity and the behavior of an MSA under ML based phylogenetic analysis.

### Machine Learning and Evaluation

During our experiments, we trained distinct regression algorithms and compared their predictive power according to the R^2^ score, the MSE, the MAE, and the MAPE. We divide the training data into two sets: a training set and a test set. We use the training set to train the prediction algorithms and the test set to evaluate the trained predictors on unseen data. We train multiple different regression models, namely Linear Regression, Lasso Regression (Tibshirani 1996), Random Forest Regression (Ho 1995), Adaptive Boosting (AdaBoost) (Freund and Schapire 1996), and Support Vector Regression (Boser *et al*. 1992). Random Forest Regression proves to be the most suitable Machine Learning algorithm for the task at hand, and outperforms all other tested regression models according to all our metrics. In Supplementary Information Section 3, we present the results for all trained regression models. Random Forest Regression is an ensemble method that averages over the predictions of multiple independently trained decision trees. To determine the optimal set of hyperparameters for the Random Forest Regression, we implemented a grid search that tests various combinations of hyperparameter values. For this grid search, we use an additional validation set, obtained by further subdividing the training set. We then perform hyperparameter optimization using this validation set. Our final difficulty predictor consists of 100 decision trees with a maximum depth of 10. To prevent overfitting, we set the minimum number of samples in a leaf node to 10 and the minimum number of samples required for a split to 20. Further, we train the individual decision trees on bootstrapped training data. We set the sample size for the bootstrapping to 75 % of the training data size. Note that this bootstrapping procedure samples the training data (features and corresponding label) and is not the phylogenetic bootstrap.

### Retraining the Model

To continuously and automatically improve the prediction accuracy of Pythia, we regularly extend the training data set and subsequently retrain the predictor. We extend the training data using the anonymized MSAs that we continuously obtain during our RAxML Grove database updates. Note that these MSAs are only available internally in RAxML Grove and are not publicly available. To limit the amount of resources required for retraining, we do not include every incoming, new MSA. We select MSAs based on a heuristic instead. At the time of writing, we select the set of new MSAs such that it diversifies the distribution of features in our training data. Algorithm 1 shows the heuristic for deciding whether to use a given MSA for retraining. For each feature *f*_*i*_, we compute the respective histogram *H*_*i*_ on the training data using a predefined number of bins *n*_bins_. Next, we compute the respective feature value for the given MSA and find the corresponding bin *hist bin* in the histogram *H*_*i*_. The goal is to attain an even distribution of features, that is, all histogram bins should have the same height 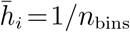. To quantify the deviation *v*_*i*_ from this even distribution, we divide this desired height 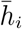 by the actual height *h*_*i*_ of *hist bin*. The deviation *v*_*i*_ is negatively correlated to the number of samples in the corresponding histogram bin. For bins with fewer samples than the desired even distribution, the deviation is > 1. We sum the deviations *v*_*i*_ across all features. We use the given MSA for retraining if this sum is ≥ 14 or any of the deviations *v*_*i*_ is ≥ 4. The rationale for the first threshold is that in this case, on average, for each feature *f*_*i*_ the corresponding bin *hist bin* has only half the desired height. The rationale for the second threshold is that in this case, one of the feature bins has only 1/4-th of the desired height.

#### Algorithm 1

Heuristic for deciding whether to use a given MSA for retraining Pythia.

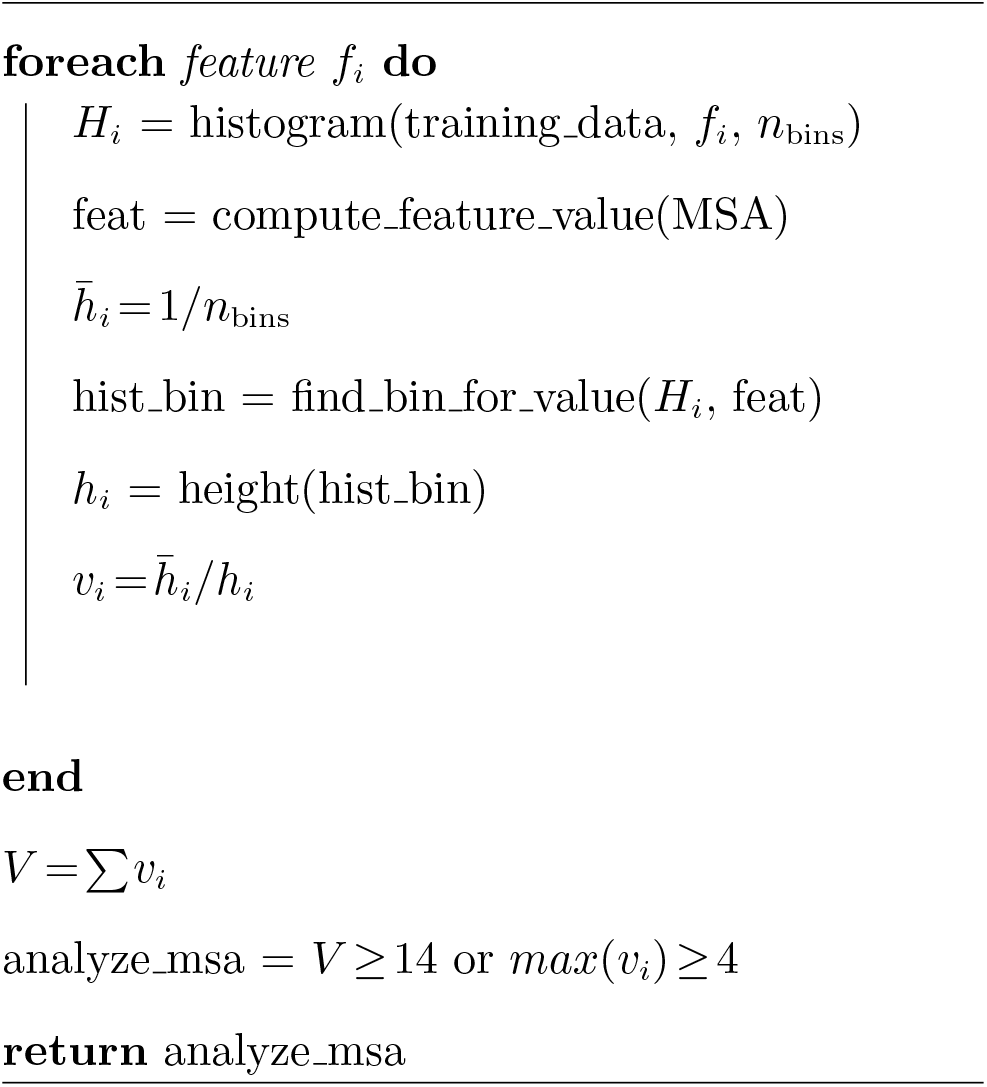

For all MSAs we select, we compute the ground-truth label and prediction features as described in the Training Data subsection. Based on this enlarged training data, we retrain Pythia and automatically update the trained predictor in our Python and C libraries.

## Code and Data availability

We provide Pythia as open source software libraries in C and Python. Both libraries include the trained Random Forest Regressor and the computation of the required prediction features. The C library CPythia is an addition to Coraxlib and is available at https://github.com/tschuelia/CPythia.

Additionally, we provide PyPythia, a lightweight, stand-alone Python library, including a command line interface. PyPythia is available at https://github.com/tschuelia/PyPythia. The implemented pipeline to compute the prediction features and ground-truth difficulty labels for the training data is available at https://github.com/tschuelia/difficulty-prediction-training-data. This repository also contains the training data as parquet file.

## Supplementary Information

Supplementary information is available online.

## Acknowledgments and Funding

The authors gratefully acknowledge the support of the Klaus Tschira Foundation. This project has received funding from the European Union’s Horizon 2020 research and innovation programme under the Marie Sklodowska-Curie grant agreement No 764840.

