## Supplementary Information for "From Easy to Hopeless - Predicting the Difficulty of Phylogenetic Analyses"

October 10, 2022

#### 1 Training Data

We trained the prediction model using empirical datasets obtained from TreeBASE [13]. To date, our training data consists of 3250 MSAs, of which 74% are DNA MSAs and 26% are protein MSAs. The training data includes partitioned and unpartitioned MSAs. Figure S1 shows the distribution of the number of taxa, sites, patterns, partitions, the proportion of gaps and invariant sites, as well as the sites-over-taxa ratio, and patterns-over-taxa ratio in the training data. Figure S2 shows the distribution of assigned difficulties according to the definition we present in the main paper.

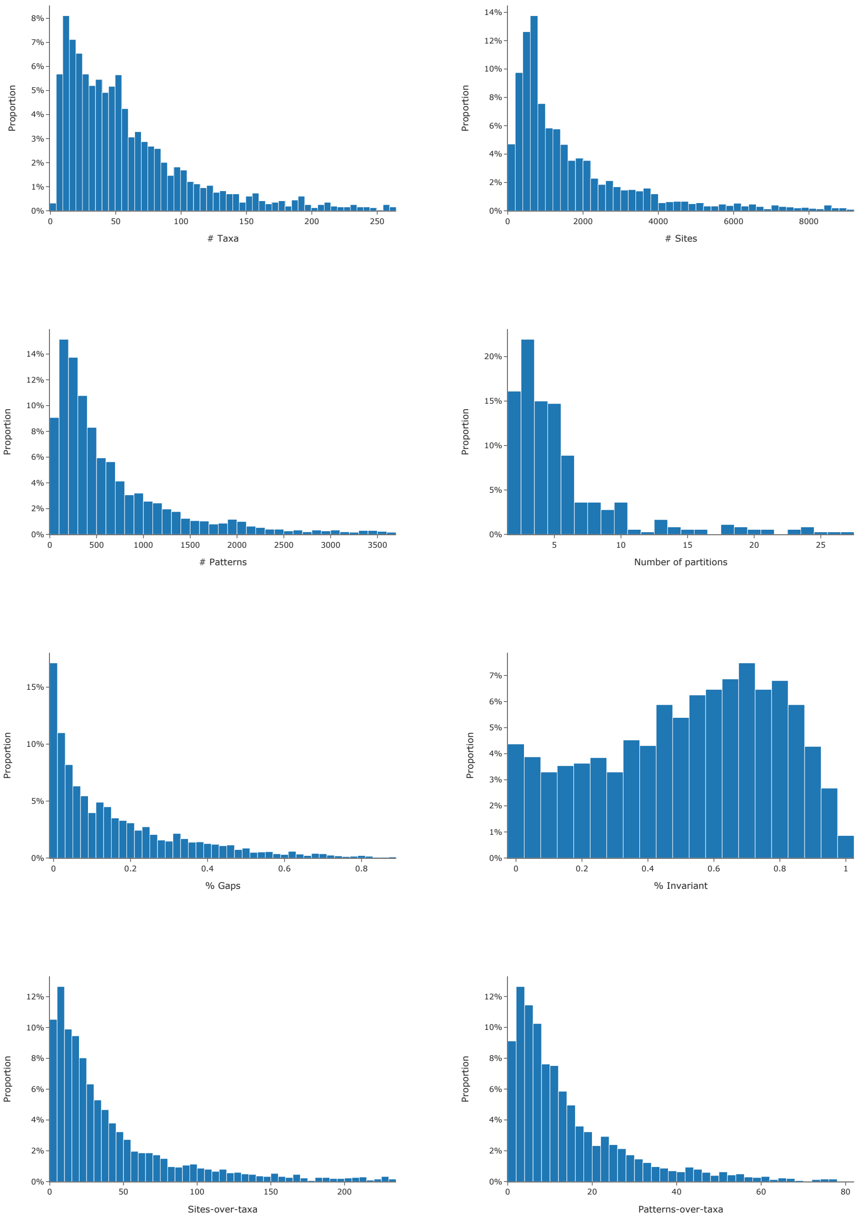


Figure S1: Distribution of MSA attributes in the training data.

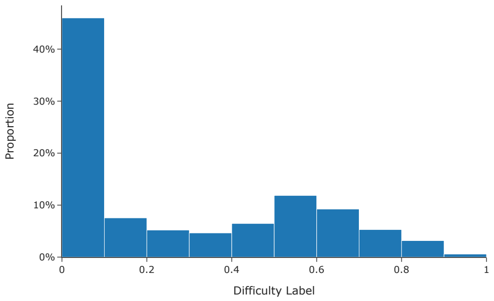


Figure S2: Distribution of assigned difficulty labels in the training data.

### 2 Feature Engineering

In our preliminary experiments, we assessed a plethora of distinct features based on the MSA, trees inferred under parsimony [3, 4], and features based on a single ML tree inference using RAxML-NG [9]. In this section, we present all features we implemented and analyzed, compare their feature importances and runtimes, and motivate the selection of the features we use.

#### 2.1 MSA Features

**Sites-over-taxa ratio**

$$\frac{\#\text{ Sites}}{\#\text{ Taxa}} = \frac{\text{Number of alignment columns}}{\text{Number of taxa}}$$

**Patterns-over-taxa ratio:**

$$\frac{\#\text{ Patterns}}{\#\text{ Taxa}} = \frac{\text{Number of unique sites}}{\text{Number of taxa}}$$

**% Invariant sites:** Percentage of fully conserved sites.

**% Gaps:** Proportion of gaps in the MSA.

**MSA Entropy** The Shannon Entropy [15] measures the information value of data. We compute the entropy for an MSA with $N$ taxa as average Shannon Entropy over all $M$ MSA sites/columns:

$$H(\text{MSA}) = \frac{1}{M}\sum_{i=1}^{M} H(\text{site}_i) \tag{1}$$

$$\text{with} \quad H(\text{site}_i) = -\sum_{j=1}^{N} P(\text{char}_{j,i}) \cdot log(P(\text{char}_{j,i})) \tag{2}$$

**Bollback Multinomial** Bollback [1] designed this test statistic to quantify the frequency of site patterns. Let $M$ be the length of the MSA, $n$ the number of unique site patterns, $\xi(i)$ the $i$-th unique pattern, and $N_{\xi(i)}$ the number of times this pattern occurs. We compute the multinomial test statistic as:

$$T(\text{MSA}) = \left(\sum_{i=1}^{n} N_{\xi(i)} \cdot ln(N_{\xi(i)})\right) - M \cdot ln(M) \tag{3}$$

**Treelikeness Score** The treelikeness score [7] is designed to quantify the phylogenetic signal in the data. The treelikeness score is based on a matrix of pairwise distances between all sequences in the MSA. Let $N$ be the number of taxa, and $D$ the pair-wise distance matrix with entries $d_{i,j}$; $0 \leq i, j \leq N$. For a set of four taxa (a *quartet*) $q = (x, y, u, v)$, we compute the treelikeness as:

$$\delta_q = \frac{d_{xv|yu} - d_{xu|yv}}{d_{xv|yu} - d_{xy|uv}} \tag{4}$$

$$\text{with} \quad d_{xy|uv} = d_{xy} + d_{uv} \tag{5}$$

$$\text{and} \quad d_{xy|uv} \leq d_{xu|yv} \leq d_{xv|yu} \tag{6}$$

The lower the score $\delta_q$, the stronger the phylogenetic signal in the quartet $q$ will be. For the set $N$ of taxa, we compute this $\delta_q$-score for every possible quartet $q$ in $N$. The treelikeness of the entire data is then computed as the mean over all $\delta_q$-scores. Due to the computational complexity, for datasets with many taxa ($> 100$), Holland *et al.* [7] suggest computing the $\delta_q$-scores for a random sample of all possible quartets.

#### 2.2 Tree Features

Our difficulty prediction aims to reduce the amount of resources used for analyzing easy MSAs, and to increase user awareness about the degree of difficulty of an MSA and its expected behavior in tree space *prior* to tree actual tree inference. However, when using the ML method for the subsequent analysis, we can enrich the set of features by inferring a single ML tree. For our analyses, we use the ML search heuristic implemented in RAxML-NG to infer this single tree. If these features prove to impact the prediction substantially, instead of predicting the difficulty prior to any tree inference, we predict the difficulty only *after* a single tree inference. This, however, induces a substantial runtime increase. We analyze the following set of features based on a single RAxML-NG tree inference:

- # SPR rounds: RAxML-NG optimizes the tree topology of an initial starting tree using Subtree Pruning and Regrafting (SPR) moves. One iteration of pruning and regrafting all possible subtrees in the current tree is called an *SPR round*. This feature counts the number of SPR rounds RAxML-NG performs during this single tree inference.
- Branch lengths: Minimum, maximum, average, standard deviation, and sum over all branch lengths in the inferred tree.
- RF-Distance: Topological distance between the starting tree topology and the inferred tree topology using the Robinson-Foulds distance metric (RF-Distance) [14].

#### 2.3 Parsimony Features

Compared to the ML method, parsimony scores can be computed orders of magnitude faster, while the underlying tree search problem is the same. In our analyses, we infer 100 trees using the parsimony-based randomized stepwise addition order heuristic implemented in RAxML-NG. Based on the inferred parsimony trees, we compute the following features:

- RF-Distance: Average pairwise relative RF-Distance between the 100 inferred trees.
- % Unique topologies: Proportion of unique tree topologies among the 100 inferred trees.

#### 2.4 Feature Importance and Runtime

As stated in the main paper, we only selected a subset of all features to train the final difficulty predictor. To select this subset of features, we first trained a Random Forest Regressor using all presented features and considered the respective feature importances, as well as the runtimes to obtain the respective features. Table S1 shows the permutation-based importance of each of the presented features in percent.

Note that the feature importances reported in Table S1 deviate from the feature importances reported in Table 1 in the main paper. We trained the final predictor of Pythia using only the subset of eight features as presented in this section, yielding the feature importances we show in Table 1 in the main paper. In contrast, the feature importances we present in Table S1 refer to a predictor that we trained using all features we initially considered for the difficulty prediction task.

| Feature | Importance |
|---|---|
| RF-Distance parsimony trees | 54.1 % |
| Total branch length | 9.3 % |
| # SPR rounds | 5.7 % |
| Entropy | 5.2 % |
| Treelikeness | 3.8 % |
| RF-Distance starting – final | 3.5 % |
| Sites-over-taxa | 3.5 % |
| Bollback | 3.1 % |
| Patterns-over-taxa | 3.0 % |
| Average branch length | 2.5 % |
| Std. dev. branch length | 1.9 % |
| % Gaps | 1.6 % |
| % Invariant | 1.0 % |
| Maximum branch length | 0.9 % |
| % Unique topologies parsimony trees | 0.7 % |
| Minimum branch length | 0.2 % |

Table S1: Permutation-based feature importance of each feature we consider for predicting the difficulty.

Table S2 shows the runtimes to compute the presented features. The table shows the average absolute runtime in seconds, as well as the average runtime relative to the runtime of a single RAxML-NG tree inference. For benchmarking the runtimes of the feature computation, we used the implementation in our Python library PyPythia, and we ran the benchmark on a Xeon Platinum 8260 CPU (48 cores with 2.4 GHz and 754 GB RAM). Since tree features require a complete ML tree search, the relative compute time for these features is $\geq$ 100 %. For obtaining the number of taxa, sites, patterns, the proportion of gaps and invariant sites, we use the RAxML-NG `--parse` option. Since RAxML-NG log files contain but a few lines, parsing the respective files using Python is fast (runtime $\ll$ 10 ms). We therefore simply omit this additional runtime in our assessment.

| Feature | Absolute Runtime in seconds $\mu \pm \sigma$ (median) | Relative runtime in percent $\mu \pm \sigma$ (median) |
|---|---|---|
| **MSA Features:** | | |
| Sites-per-taxa, patterns-per-taxa, % gaps, % invariant | $0 \pm 0\,\mathrm{s}$ (0 s) | $1.1 \pm 3.1\,\%$ (0.2 %) |
| Entropy | $1 \pm 5\,\mathrm{s}$ (0 s) | $8.8 \pm 65.8\,\%$ (1.5 %) |
| Bollback multinomial | $0 \pm 2\,\mathrm{s}$ (0 s) | $3.3 \pm 16.6\,\%$ (0.9 %) |
| Treelikeness | $96 \pm 337\,\mathrm{s}$ (27 s) | $450.5 \pm 965.0\,\%$ (254.3 %) |
| **Tree Features:** | | |
| Single tree inference (includes the # SPR rounds) | $393 \pm 2265\,\mathrm{s}$ (12 s) | $100.0 \pm 0.0\,\%$ (100.0 %) |
| RF-Distance starting − final | $393 \pm 2265\,\mathrm{s}$ (12 s) | $100.8 \pm 2.5\,\%$ (100.1 %) |
| Branch length statistics | $393 \pm 2265\,\mathrm{s}$ (12 s) | $100.1 \pm 0.2\,\%$ (100.0 %) |
| **Maximum Parsimony Features:** | | |
| 100 parsimony trees, RF-Distance, % unique topologies | $4 \pm 28\,\mathrm{s}$ (0 s) | $8.2 \pm 13.4\,\%$ (3.8 %) |

Table S2: Runtime to compute the features we consider for difficulty prediction.

Our goal is to train a prediction algorithm that is both accurate and fast. We therefore select a subset of the presented features with a good trade-off between accuracy and runtime. In particular, omitting the tree features and the treelikeness score improves the prediction speed substantially. Thus, despite the high feature importances of the tree features, we decided against using them. By relying on the subset, accuracy only decreases slightly, as Table S3 shows.

| Features | $R^2$ Score | MSE | MAE | MAPE |
|---|---|---|---|---|
| All features | 0.81 | 0.01 | 0.08 | 2.1 % |
| Subset | 0.79 | 0.02 | 0.09 | 2.9 % |

Table S3: Prediction accuracy when using all features, compared to when using only the selected subset of features.

The runtime overhead of computing the features depends predominantly on the MSA size. Figure S3 shows the runtime to compute the selected subset of features relative to the runtime for a single tree inference, depending on the MSA size. For better visualization, we filtered MSA size outliers using Tukey's fences [21] with $k = 3$ prior to plotting.

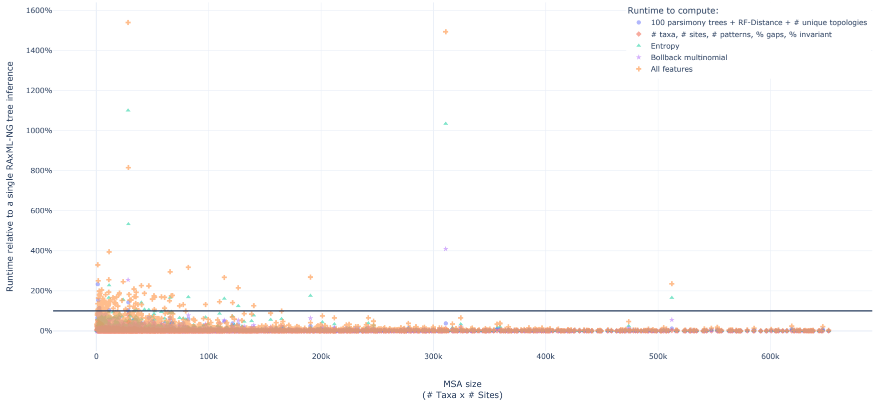


Figure S3: Runtimes to compute the subset of features used for predicting the difficulty. The x-axis states the size of the MSA (# Taxa · # Sites) and the y-axis the runtime relative to the runtime for inferring a single tree using RAxML-NG. The black horizontal line denotes $y = 100\,\%$.

### 3 Regression Algorithms

#### 3.1 Comparison of Algorithms

During our initial experiments, we trained and compared different machine learning regression algorithms. We use the implementation of the regression algorithms of the Python machine learning library scikit-learn [12]. Table S4 shows the $R^2$ score, mean squared error (MSE), mean absolute error (MAE) and mean absolute percentage error (MAPE) of all trained regressors. Additionally, we use dummy regressors as baseline. These dummy regressors do not learn to distinguish the data, but predict a value based on predefined rules. We use three dummy regressors: one that always predicts the mean difficulty of the training set, one that always predicts the 0.75 quantile of the training set, and one that predicts 0.5 for each dataset. The table shows that the Random Forest Regressor outperforms all other trained regressors according to all metrics. Using the dummy regressors as baseline, we observe that the trained regressors (except for the Support Vector Regression) learned to distinguish datasets according to their difficulty.

| Algorithm | $R^2$ Score | MSE | MAE | MAPE |
|---|---|---|---|---|
| Random Forest Regressor [6] | **0.79** | **0.02** | **0.09** | **2.9 %** |
| Linear Regression | 0.66 | 0.03 | 0.14 | 7.2 % |
| AdaBoost [5] | 0.64 | 0.03 | 0.15 | 9.7 % |
| Lasso Regression [20] | 0.01 | 0.08 | 0.26 | 18.0 % |
| Dummy Regressor (mean) | -0.0 | 0.08 | 0.26 | 18.3 % |
| Support Vector Regression [2] | -0.06 | 0.09 | 0.25 | 12.1 % |
| Dummy Regressor (constant) | -0.6 | 0.13 | 0.32 | 33.7 % |
| Dummy Regressor (quantile) | -0.84 | 0.15 | 0.34 | 36.4 % |

Table S4: Results of the trained predictors. The bold values indicate the highest scoring regressor per metric.

### 4 Training Error

As stated in the main paper, we observe that Pythia tends to overestimate the difficulty of easy MSAs and tends to underestimate intermediate and difficult datasets. As Figure S4 depicts, this observation holds true for both, the training, and the test set.

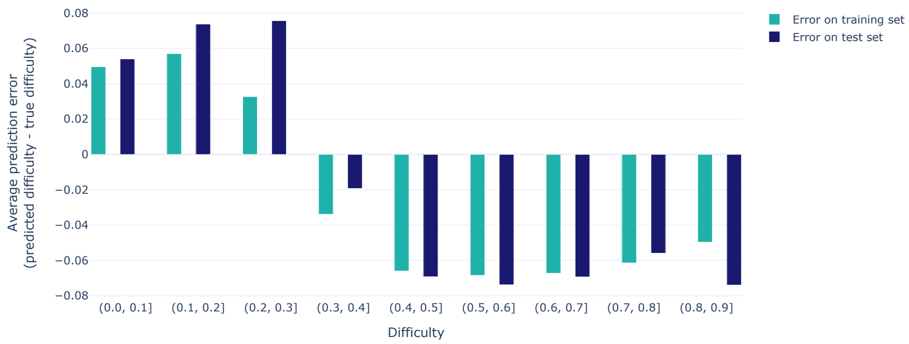


Figure S4: Average prediction error per difficulty range for the training and test sets. The prediction error is computed as *predicted difficulty - true difficulty*.

Figure S5 shows the distribution of prediction error for the training data. Figure S5a shows the absolute prediction error and Figure S5b shows the relative prediction error. Both figures show the errors for the training and the test set separately.

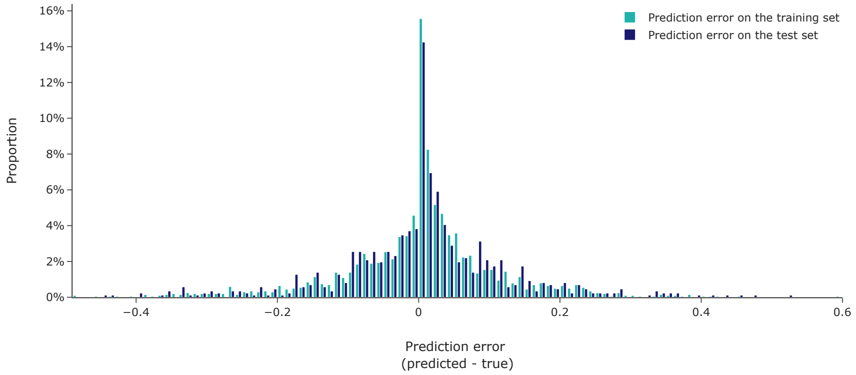


(a) Distribution of the absolute prediction error.

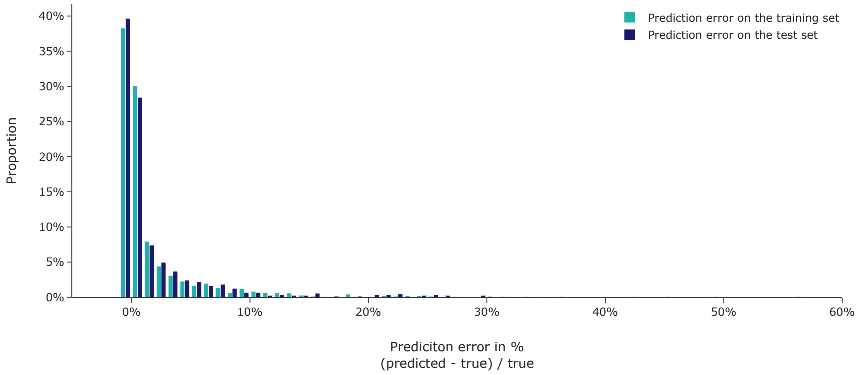


(b) Distribution of the prediction error in percent.

Figure S5: Distribution of prediction errors for the training and test set.

### 5 DNA and AA data

In our analyses, we do not explicitly distinguish between DNA and AA data, as Pythia can predict the difficulty for both data types. DNA data accounts for 74% of our training data and AA data for 26%. To better justify the equal treatment of DNA and AA data, we analyzed the distribution of feature values for all eight prediction features for DNA and AA data separately. Figure S6 shows the distribution of the number of taxa, sites, patterns, the proportion of gaps and invariant sites, as well as the sites-over-taxa ratio, and patterns-over-taxa ratio, entropy and the bollback multinomial on a per data type basis. Figure S7 depicts the distribution of assigned difficulties according to the definition we present in the main paper.

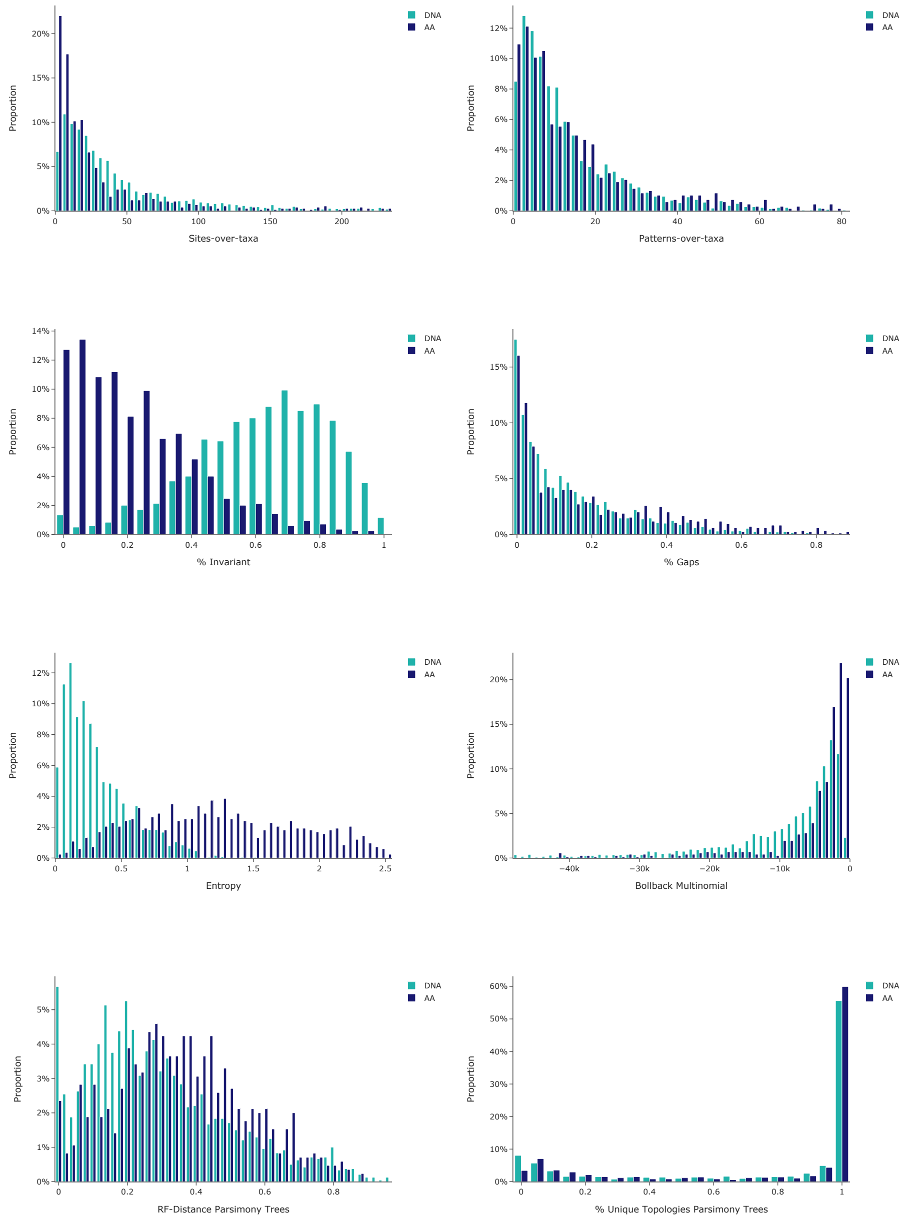


Figure S6: Distribution of prediction feature values in the training data, separated by data type (DNA and AA).

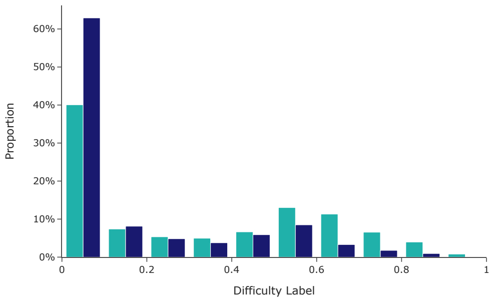


Figure S7: Distribution of assigned difficulty labels in the training data, separated by data type (DNA and AA).

Across all eight features we use for difficulty prediction with Pythia, DNA data and AA data behave in an almost identical manner. We do, however, observe a noticeable difference in the proportion of invariant sites and the entropy. While the proportion of invariant sites has a very low feature importance (0.6%, see Table 1 in the main paper), the entropy has a high feature importance (17%). The mean and standard deviation of the entropy for DNA data is $\mu \pm \sigma = 0.32 \pm 0.25$, whereas for AA data it is $1.26 \pm 0.64$. However, when further analyzing the prediction error of Pythia, we notice that the distribution of prediction errors is similar for DNA and AA data, indicating that Pythia has a similar prediction accuracy for both, DNA, and AA data (Figure S8). We conclude that including DNA and AA in the same predictor does not influence the prediction accuracy for either data type, and that a further distinction is not necessary.

| Tool | Software Version |
|---|---|
| RAxML-NG | Adapted version based on version 1.1.0. Available at `https://github.com/tschuelia/raxml-ng`. |
| IQ-TREE | Version 2.1.3, available at `https://github.com/iqtree/iqtree2/releases/tag/v2.1.3` |

Table S5: The software versions of RAxML-NG and IQ-TREE we used for our analyses.

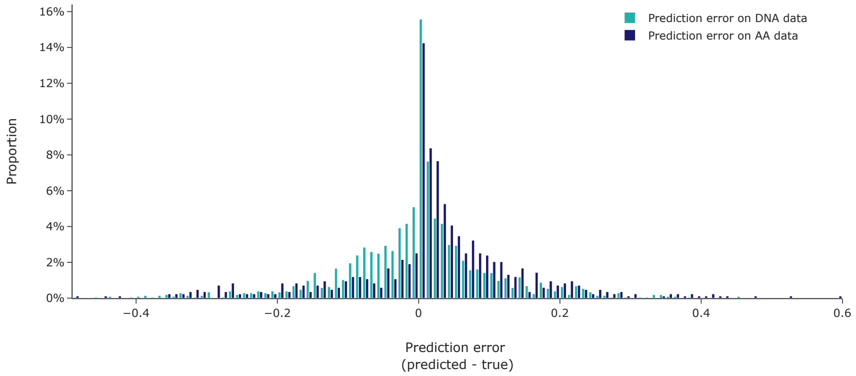


Figure S8: Distribution of the absolute prediction error on the training data, separated by data type (DNA and AA). The prediction error is computed as *predicted difficulty - true difficulty*.

### 6 Software and Command Lines

In our analyses, we use RAxML-NG to infer Maximum Likelihood trees. We first infer a tree using the standard tree inference mode, and afterwards re-evaluate the log-likelihood using the `--eval` mode. In this execution mode, RAxML-NG does not alter the tree topology, but only optimizes the branch lengths and substitution model parameters. We use the significance tests implemented in IQ-TREE [11] to determine the plausible tree sets. Table S5 states the software versions of RAxML-NG and IQ-TREE we use. As the table states, we use an adapted version of RAxML-NG. This version logs the parsimony scores of the parsimony trees that RAxML-NG uses as initial starting tree topologies. Additionally, in this version, RAxML-NG clusters input trees based on the RF-Distance when run in `--rfdist` mode. Trees within a cluster have identical tree topologies (RF-Distance = 0.0). Using this clustering, we filter duplicate tree topologies prior to the statistical significance testing.

IQ-TREE implements the following significance tests: the Kishino-Hasegawa test [8] and the Shimodaira-Hasegawa test [16], both in their weighted and unweighted variants, the Approximately Unbiased test [17], as well as the Expected Likelihood Weight test [18]. We use the default IQ-TREE settings for the number of resampling of estimated log-likelihoods (RELL) replicates (10 000) and the significance level ($\alpha = 0.05$). Since the significance tests can be biased by the number of trees in the candidate set [18], we remove topologically identical trees before applying the tests.

For unpartitioned DNA MSAs we use the general time reversible (GTR) model [19] of nucleotide substitution as it is the most flexible and general model for nucleotide substitution. We use four discrete $\Gamma$ rate categories to account for among site rate heterogeneity. We use the LG substitution model [10] with four discrete $\Gamma$ rate categories for unpartitioned AA MSAs.
